# Remembering the pattern: A longitudinal case study on statistical learning in spatial navigation and memory consolidation

**DOI:** 10.1101/2021.10.18.464818

**Authors:** Kathryn N. Graves, Brynn E. Sherman, David Huberdeau, Eyiyemisi Damisah, Imran H. Quraishi, Nicholas B. Turk-Browne

**Affiliations:** Department of Psychology, Yale University, 2 Hillhouse Ave., New Haven, CT 06520, USA; Department of Neurosurgery, Yale University, 333 Cedar St., New Haven, CT 06510, USA; Department of Neurology, Yale University, 800 Howard Ave., New Haven, CT 06519, USA; Wu Tsai Institute, Yale University, 100 College St, New Haven, CT 06510, USA

**Keywords:** memory integration, complementary learning systems, intracranial EEG, hippocampus, medial prefrontal cortex, inverted encoding models

## Abstract

Distinct brain systems are thought to support statistical learning over different timescales. Regularities encountered during online perceptual experience can be acquired rapidly by the hippocampus. Further processing during offline consolidation can establish these regularities gradually in cortical regions, including the medial prefrontal cortex (mPFC). These mechanisms of statistical learning may be critical during spatial navigation, for which knowledge of the structure of an environment can facilitate future behavior. Rapid acquisition and prolonged retention of regularities have been investigated in isolation, but how they interact in the context of spatial navigation is unknown. We had the rare opportunity to study the brain systems underlying both rapid and gradual timescales of statistical learning using intracranial electroencephalography (iEEG) longitudinally in the same patient over a period of three weeks. As hypothesized, spatial patterns were represented in the hippocampus but not mPFC for up to one week after statistical learning and then represented in the mPFC but not hippocampus two and three weeks after statistical learning. Taken together, these findings clarify that the hippocampus may do the initial work of extracting regularities and transfer these integrated memories to cortex, rather than only storing individual experiences and leaving it up to cortex to extract regularities.

**Highlights:**

- Case study of an epilepsy patient tested longitudinally over three weeks.
- We tracked time-dependent changes in neural representations of spatial patterns.
- Representations reconstructed from hippocampal activity reflected patterns learned within a week.
- Representations reconstructed from activity in the mPFC reflected patterns learned 2–3 weeks ago.

## 1. Introduction

When you move to a new city, even on your first walk around town, you begin mapping out the streets and landmarks in your head, allowing you to subsequently find your way home that day. This knowledge grows over time through continued experience, such that it can help you find particular places on future excursions that you have never visited before. This ability to immediately represent the structure of one’s surroundings and to retain and refine this knowledge over time reflects two different timescales of *statistical learning*, or extraction of regularities from the environment.

On rapid timescales of minutes to hours, statistical learning occurs robustly across modalities (Sherman, Graves, & Turk-Browne, 2020), from auditory sequences of tones (Saffran, Johnson, Aslin, & Newport, 1999) to visual series of shapes (Turk-Browne, Jungé, & Scholl, 2005) to haptic properties (Lengyel et al., 2019), and within multiple domains, from abstract categories (Brady & Oliva, 2008) to music (Leung & Dean, 2018) to faces (Dotsch, Hassin, & Todorov, 2017). It supports parsing of streams of speech into words (Karuza et al., 2013; McNealy, Mazziotta, & Dapretto, 2006), as well as learning of motor sequences (Janacsek et al., 2020) and reward contingencies (Goldfarb, Chun, & Phelps, 2016). In the brain, this process relies at least in part on the hippocampus (Schapiro & Turk-Browne, 2015), which computational models have shown contains the necessary architecture for rapid, online learning of regularities from the environment (Schapiro, Turk-Browne, Botvinick, & Norman, 2017).

On more gradual timescales of days to weeks, or even years, cortical consolidation of encoded experiences shapes our semantic knowledge, supporting the emergence of spatial, contextual, and conceptual schemas (Gilboa & Marlatte, 2017). The formation of schemas is supported by the medial prefrontal cortex (mPFC), which represents overlapping features of previously encoded stimuli in humans (Tompary & Davachi, 2017) and spatial locations in rodents (Richards et al., 2014). For example, this latter study established a causal link between rodent mPFC and pattern consolidation by having rats learn locations in a Morris Water Maze that were drawn from an underlying, hidden distribution. After learning the individual locations, the mPFC of rats in the experimental group was disabled via a pharmacological manipulation, whereas the mPFC of rats in the control group was left intact. During a test phase 30 days later, the rats were placed back in the water maze to measure whether they had extracted the distribution through consolidation of individual locations. Rats with an intact mPFC searched according to the distribution, indicating statistical learning, but those with a impaired mPFC did not. This finding suggested a critical role for mPFC in consolidation and prolonged retention of spatial regularities.

It remains unclear how the multiple timescales of statistical learning relate to spatial navigation in humans. For example, whereas it takes a month for rodents to extract spatial patterns during navigation, humans can do so immediately during online behavior (Graves, Antony, & Turk-Browne, 2020). Participants in this study performed a virtual analogue of the Morris Water Maze, finding hidden locations in an arena that were drawn from a Gaussian distribution. Over the course of a few minutes of learning, navigation behavior switched from a bias toward previously learned individual locations to a bias toward the mean of the distribution of locations.

Despite being acquired rapidly, the fate of this acquired representation over time is unknown. Moreover, whereas we hypothesize that the hippocampus may support rapid extraction, followed by more prolonged retention in mPFC, such time-dependent changes in the brain systems involved in human navigation have not been tested previously.

In the current study, we tracked the cognitive and neural trajectory of learned spatial patterns during navigation via intracranial electroencephalography (iEEG). We had the rare opportunity to repeatedly test a single patient, who was in the hospital for clinical seizure monitoring substantially longer than an average study (6 weeks vs. 1-1.5 weeks typically). The patient performed a screen-based virtual maze task (Graves et al., 2020) where hidden target locations were drawn from a Gaussian distribution. She learned two distributions of locations on two separate days of training, after which the extent of her distribution extraction was tested at multiple intervals. We predicted that the hippocampus would represent the underlying distribution initially after learning, but that this pattern would come to be represented in mPFC over time.

## 2. Materials and Methods

### 2.1 Participant

We tested one patient (female, age 26, right-handed) admitted to Yale New Haven Hospital for iEEG seizure monitoring. She had a history of epilepsy and attention deficit hyperactivity disorder (ADHD) starting at age 10, as well as recurring depression starting when she was a teenager. Her medications included oxcarbazepine and intronasal midazolam for epilepsy, as well as occasional Adderall for ADHD. A structural MRI in 2017 revealed normal hippocampal volume. Neuropsychological test scores were high overall (Full Scale Intelligence Quotient (FSIQ) = 131, Verbal Comprehension Index (VCI) = 141, Perceptual Reasoning Index (PRI) = 123) and non-indicative of any seizure lateralization or localization. Pre-implant video and EEG monitoring suggested seizures were likely arising from the left posterior lateral temporal region.

The patient was implanted with subdural electrodes on the cortical surface over the left hemisphere, as well as multiple depth electrodes implanted in subcortical structures including the left hippocampus (Fig. 1). Decisions on electrode placement were determined solely by the clinical care team to optimize localization of seizure foci. The patient completed three sessions of a virtual Morris Water Maze task, referred to hereafter as Day 1, Day 8, and Day 21. The research protocol was approved by the Yale University Institutional Review Board.

**Fig. 1.**
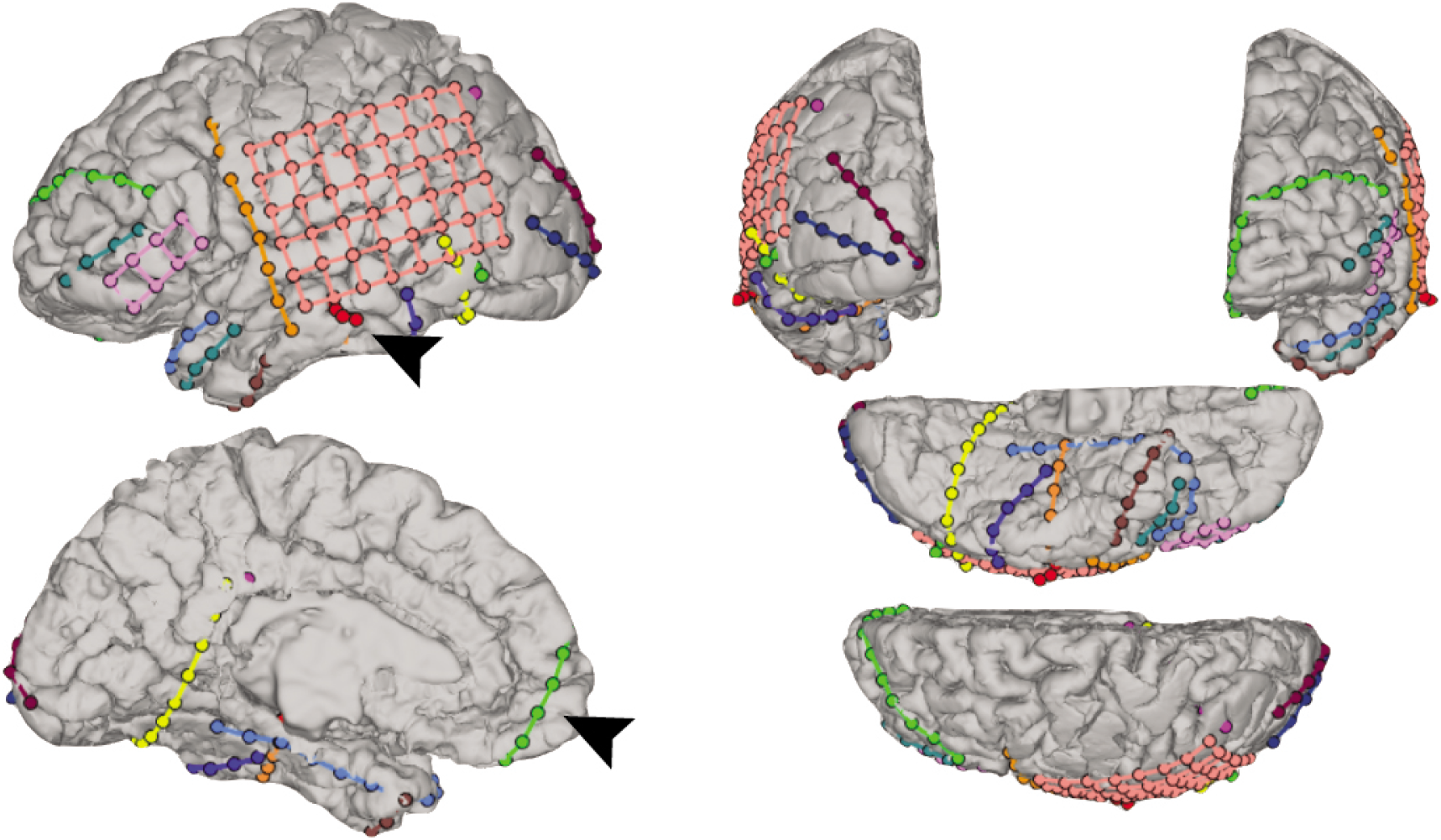
Reconstruction of the patient’s electrode coverage. Arrows are pointing to a (red) depth electrode containing left hippocampal contacts at the tip and to a (green) electrode strip covering left mPFC.

### 2.2 Experimental Design

The virtual environment was designed as a circular arena and constructed in Blender (www.blender.org). The environment was rendered in Matlab (Mathworks, Natick, MA, USA), and Psychtoolbox (Brainard, 1997) was used to display task instructions. The arena was designed graphically with an island beach theme. The circular floor (radius = 7.85 arbitrary units, AU) was covered by an image of sand. Mountains appeared on the north end and palm trees on the south end; similar to the classic water maze task, these landmarks served as directional headings. Each “platform” was a shell in the first session and a crab in the second session. Following Richards et al. (2014), the locations of the shells were drawn from a normal distribution of distances *d* from center (*μ* = 3.4 AU, *σ* = 0.9067 AU) and a circular normal distribution of angles *θ* between the platform and the eastern cardinal direction (*μ* = 0.2618 radians, *κ* = 8). The locations of the crabs were generated by taking the shell distribution and rotating it 160 degrees counterclockwise around the arena. We chose 160 degrees instead of a full 180 degree rotation to prevent the use of symmetry-based heuristics.

### 2.3 Procedure

On Day 1, the patient was trained on the shell distribution. She was instructed that she was on an island, searching for a total of 20 seashells buried below the sand. She could only find one shell at a time and had to walk around the beach until she found it. The patient used the *I* key to walk forward, the *J* and *L* keys to turn left and right, respectively, and the *M* key to walk backward. At the beginning of the search for each new shell, the patient’s location was initialized to the center of the arena, always facing a new random direction. Navigation was initiated by the patient following a minimum of 4 seconds in which the ground was green and the *I* /*M* keys were locked only allowing the patient to rotate in place left or right. The ground then turned to sand, cuing the patient that she could now start moving forward and backward. There was no time limit, and thus the searches varied in length. The patient successfully located a shell when she walked within a 1 1/3-unit radius of the platform location of that shell. When this occurred, further navigation was locked, and the walls turned green to reveal the outline of a shell. The screen then went black, and the patient was told how many new shells remained to be found before beginning to search for the next shell.

On Day 8, the patient was first trained on the crab distribution. This second distribution was introduced to provide another set of lags between training and test, over which we could examine time-dependent changes in hippocampal and mPFC representations. The crab training on Day 8 was nearly identical to the shell training on Day 1, with the only difference being that the patient was now looking for crabs instead of shells and the distribution was centered in a new part of the arena. After finding all of the crabs, the patient then completed a test phase following a brief delay for instructions. During the test phase, the patient was instructed that she was back on the island and that she would alternate between finding more shells and crabs. She received a cue on each trial indicating which object was the target. The test phase differed critically from training in that search was limited to 16 seconds and no feedback was given until the end of the time. This resulted in the patient searching an empty arena based on what was learned during training. The patient completed four alternating trials of searching for shells and searching for crabs. The Day 8 shell test evaluates a lag of 7 days, whereas the Day 8 crab test evaluates an immediate lag of 0 days.

On Day 21, the patient completed six more alternating tests of searching for shells and crabs in the empty arena (Fig. 2). The Day 21 shell test evaluates a lag of 20 days, whereas the crab test evaluates a lag of 13 days. Thus, across distributions we tested for the representation of spatial patterns 0, 7, 13, and 20 days after training.

**Fig. 2.**
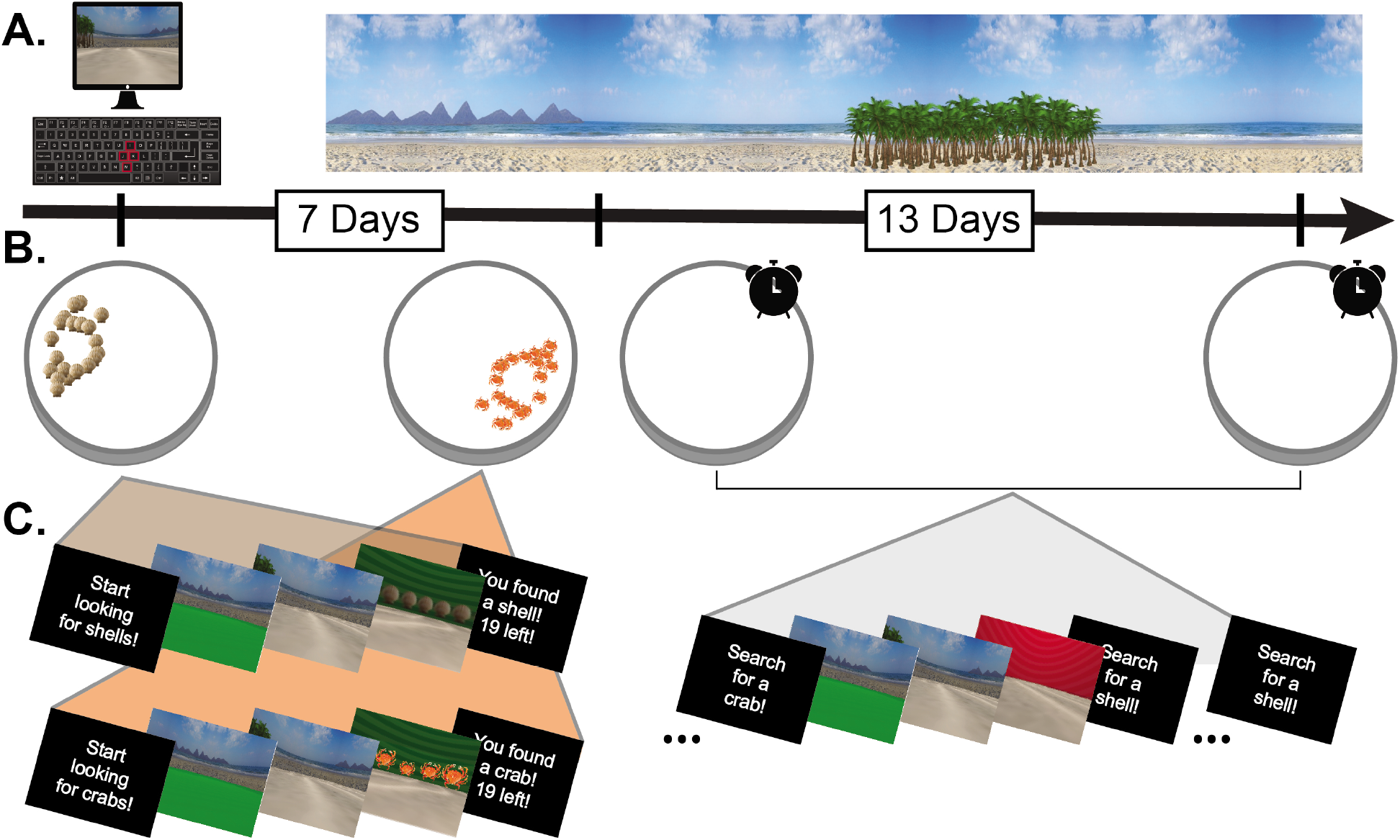
The patient’s testing schedule. A) The experiment was conducted in a virtual beach environment. B) On Day 1, the patient found a distribution of shells (brown icons). On Day 8, she found a distribution of crabs (orange icons), and then performed the first test phase, where she alternated searching for shells (7-day lag) and crabs (0-day lag) in an empty arena. She completed another alternating test phase on Day 21 for shells (20-day lag) and crabs (13-day lag). C) Each trial of training and test consisted of a cue, a pre-navigation period where forward motion was locked, a navigation period, and a post-navigation period.

### 2.4 Intracranial Recordings

Intracranial EEG (iEEG) was recorded in Natus Neuroworks 8.5.1 using a Natus Quantum amplifer, sampled at 4,096 Hz. Signals were referenced to an inverted (facing away from brain) left parieto-temporal strip electrode. In post-processing, to reduce electrical line noise, a notch filter was applied between 55-65 Hz. Data corresponding to each trial were segmented into fixation, pre-navigation, navigation, and post-navigation epochs, and downsampled to 256 Hz. To eliminate events containing epileptiform activity, epochs were removed from analysis if kurtosis of the voltage trace within epoch exceeded a threshold of 5 (van Vugt, Schulze-Bonhage, Litt, Brandt, & Kahana, 2010). This resulted in an exclusion 2.8% of experimental events. Data from the pre-navigation and navigation epochs were used for analysis.

### 2.5 Electrode Localization

Patient electrodes were localized via BioImage Suite (Papademetris, Jackowski, Rajeevan, Constable, & Staib, n.d.) and Matlab. The patient’s skull-stripped T1 weighted structural brain MRI was registered to an MNI T1 average structural MRI via nonlinear registration. The resulting transformation file was used to transform the patient’s electrode locations into standard space. Hippocampus and mPFC regions of interest (ROIs) were constructed from the Automated Anatomical Labeling (AAL) map in MNI space (Tzourio-Mazoyer et al., 2002) and used to determine which electrodes fell within each ROI.

### 2.6 Behavioral Analyses

Raw behavioral data consisted of X and Y coordinates output every 40 ms of navigation. First, we sought to confirm that by learning each hidden location during training, the patient also learning the underlying distribution. We predicted that acquisition of the spatial distribution would result in subsequent navigation that was biased towards the distribution’s central tendency.

We analyzed the extent to which learning the underlying distribution biased search behavior during the test phases by measuring whether the patient’s search path brought them closer to the location of each distribution’s mean than what would be expected by chance (Graves et al., 2020). To this end, we calculated the patient’s average proximity to the distribution mean across all test trials, separately for each distribution and testing day. We then generated a null distribution of proximity measures by rotating the spatial distribution locations a random angle between 5 and 355 degrees away from the learned distribution 1,000 times. Per rotation, we calculated the average proximity to the dummy rotated mean. An empirical *p*-value was calculated as the proportion of rotated proximity measures that were closer to the mean than the true proximity. We repeated this analysis for the distribution randomly rotated within plus and minus 90 degrees of the true distribution, and plus and minus 45 degrees, to compare search behavior to more conservative null distributions. Th objective of this analysis was to quantify the specificity of the patient’s distribution memory.

### 2.7 Time-Frequency Decomposition

We calculated a continuous Morlet wavelet transform (wave number 5) at 50 logarithmically spaced frequencies between 1 and 120 Hz for each epoch of each training and test session. A 5-second buffer was added to both ends of all power calculations to account for edge effects, and subsequently was discarded. We then estimated the background power spectrum via a linear regression fit to the power spectrum in log-log coordinates, and subtracted the subsequent 1/f from our data. Lastly, the data were z-scored per frequency band, electrode, and session. These data were used in a subsequent inverted encoding model.

### 2.8 Inverted Encoding Model

Spatial representations during test were assessed via a 2-D inverted encoding model method (Sprague & Serences, 2013). Briefly, this method enables reconstruction of a spatial representation from neural data that is modeled as a sum of the weighted activations of a set of information channels. The information channels in this study were initialized as 100 2-D Gaussians distributed throughout the surface of the beach arena, representing hypothesized place-coding activity (Fig. 3).

**Fig. 3.**
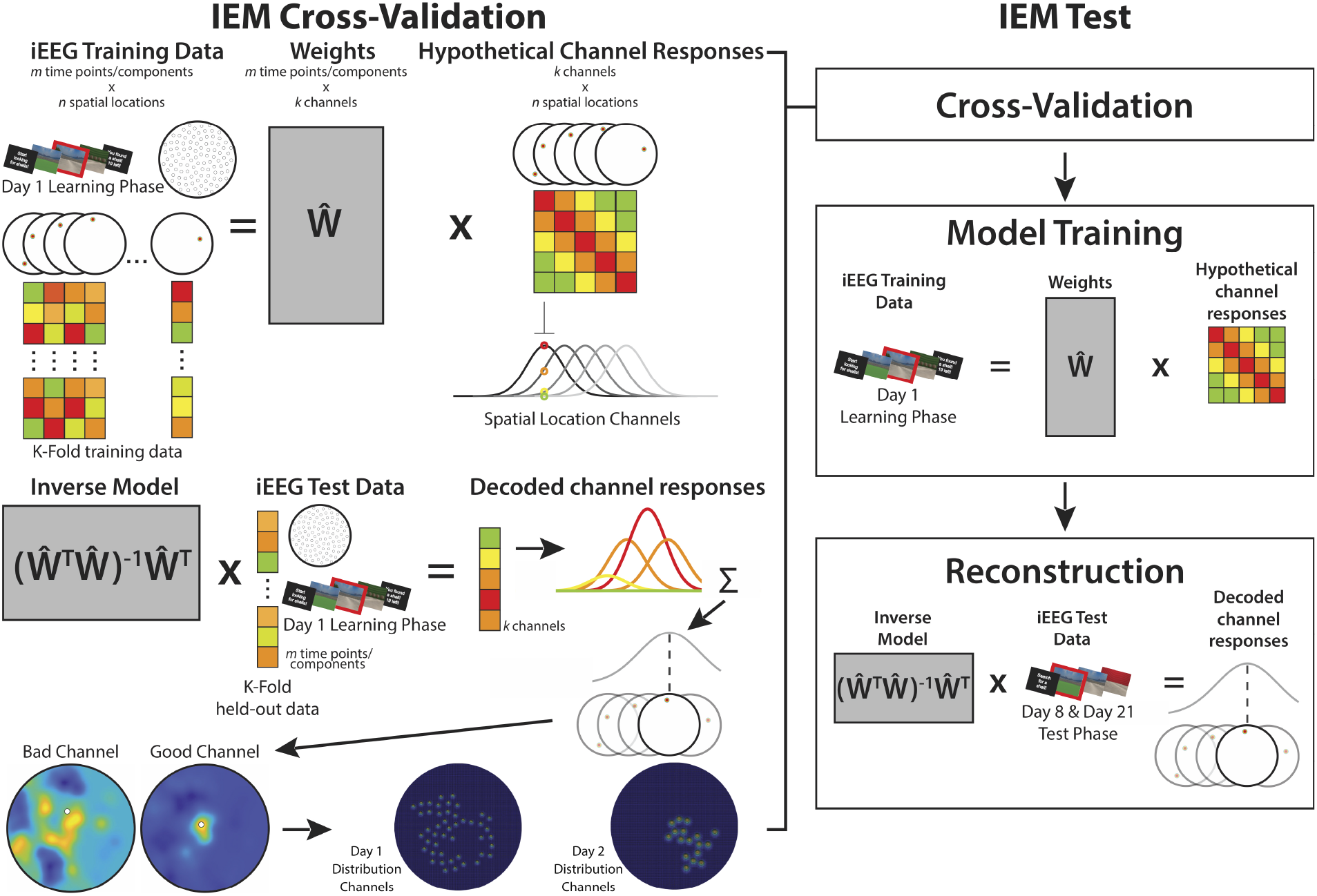
Model channel validation and test. Information channels for this model were generated based on a predicted Gaussian-shaped place field response. The model was initialized with 100 distributed channels, and navigation data from the two distribution training phases was used to train the model. The final model consisted of a subset of channels that demonstrated significant place coding activity in the training data.

**Fig. 4.**
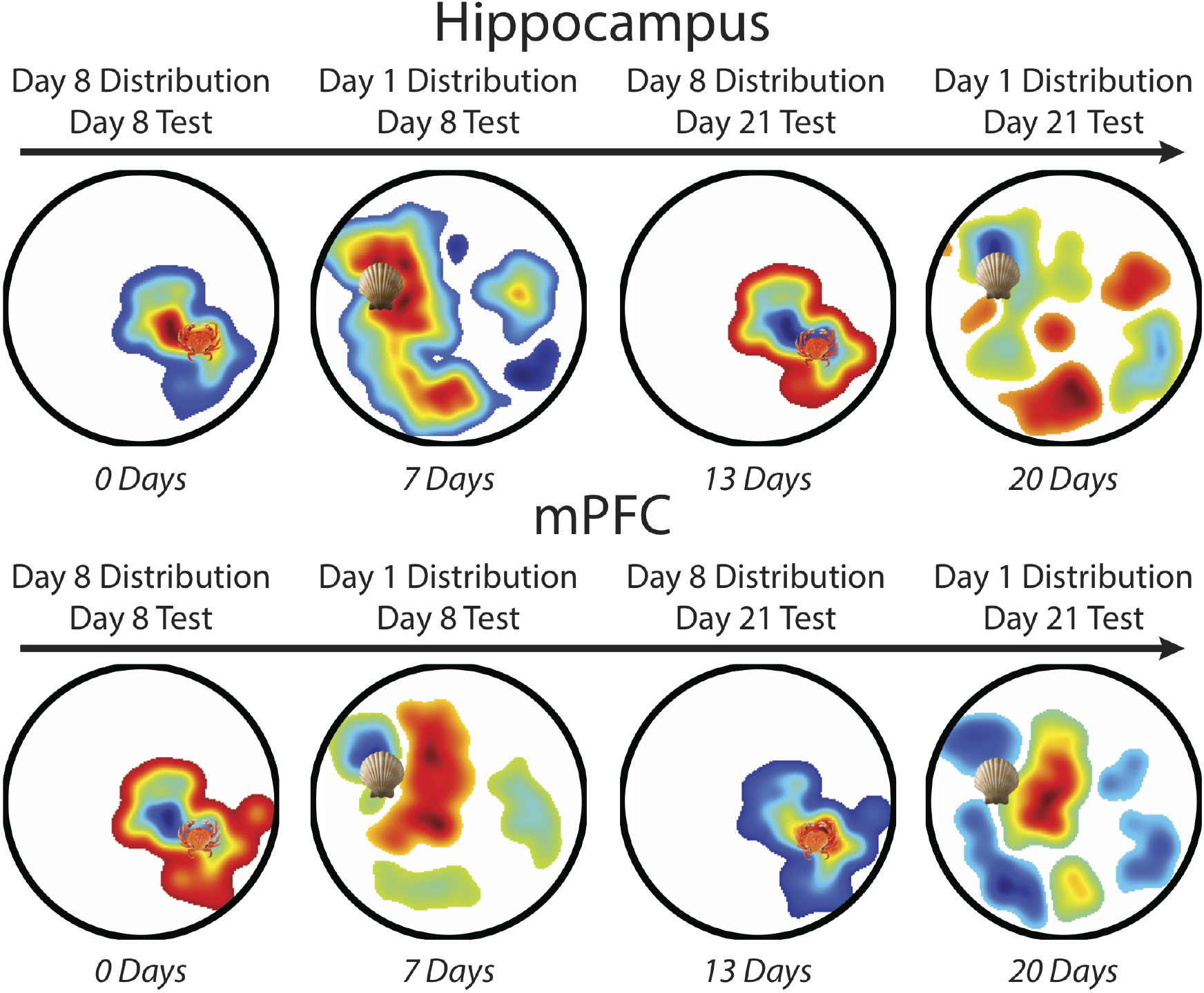
Place representations reconstructed from pre-navigation test data, plotted left-to-right as a function of lag between training and test. Each image is thresholded to exclude reconstruction from locations with no active channel. Colors are scaled by from negative to positive channel evidence, where cooler colors indicate negative channel evidence and warmer colors indicate positive channel evidence. Shell and crab icons mark the mean location from each distribution. At training/test lags of 0 and 7 days, hippocampal reconstructions were significantly proximal to the mean, but mPFC reconstructions were not. At lags of 13 and 20 days, mPFC reconstructions were significantly proximal to the mean, but hippocampal reconstructions were not.

Because the patient did not explore the entirety of the arena when learning either distribution, and therefore would not have provided neural data to train all of the channels, we conducted a channel validation step that allowed us to select for our final model only the channels for which there was enough data and that subsequently showed significant place-coding activity. This step only used data from the training phases. The resultant models were then used to reconstruct representations from unseen test phase data.

In separate models for each training day’s distribution, channel validation consisted of holding out a subset of hippocampal activity from when the patient was standing within each channel’s radius, training the model, and then testing it on the held-out training data. A channel for which peak reconstructed activity (i.e., the magnitude of the reconstructed response above the 99^th^ percentile) was more proximal to the tested channel than all other channels was considered a “good” channel (Fig. 3).

We conducted separate channel validations using time-frequency decomposed power from the 1-3 Hz band (delta, which some have termed “low theta”) and the 3-10 Hz band (theta/low alpha, which some have termed “high theta”) as model training data. Previous work in human virtual navigation has implicated hippocampal oscillations at around 3 Hz (Goyal et al., 2020; Watrous et al., 2013) and 8 Hz (Miller et al., 2018) in spatial navigation and spatial memory encoding. We thus sought to determine which band would yield greater place coding activity and recover a greater number of channels. We conducted a principal components analysis (PCA) over the frequencies by hippocampal electrodes for each band (12 frequencies between 1-3 Hz, 12 frequencies between 3-10 Hz), and iteratively repeated the channel validation on each number of components. We additionally iterated through a range of stimulus and receptive field size parameters, which determine the shape of the Gaussians (Sprague & Serences, 2013). The maximum number of recovered channels and their associated stimulus size, frequency band, and number of PCA components per distribution across all iterations served as our basis set for reconstructing the spatial representations from test data. Critically, these model optimizations were performed entirely within the training data; test data were held out of all of these steps to avoid double-dipping. Because our objective at test was to reconstruct not individual place representations, but an aggregate distribution representation, we increased the receptive field size for each distribution’s set of channels. This allowed us to generate smooth reconstructions via a similar manual parameter selection procedure as has been shown previously (Sprague & Serences, 2013).

Our main analyses build from this method of parameter and channel selection. However, the statistical trends in our results are the same if we simply set a data threshold for inclusion for all channels without establishing significant place coding activity and manually select the parameters for Gaussian shape (1 1/3 AU stimulus size, 1.2 AU receptive field size) (Supplemental Fig. 1).

To examine representations at test, we built separate models on training data from the shell and crab distributions. For each test trial, we applied the model for the corresponding cued distribution to reconstruct place activity. We used test data from only the first four pre-navigation seconds of each trial when the patient was confined to the origin of the arena, predicting that the patient would be representing whatever she had learned during training in planning her search. We combined reconstructions over all trials from a given distribution and test session, yielding a reconstruction for each lag and brain region.

We then assessed whether the peak reconstructed activity was significantly proximal to the distribution mean to quantify neural evidence of statistical learning of the distribution. We first determined the most active channel and then calculated the distance from that channel to the distribution mean. We shuffled the full reconstruction matrix, determined the most active channel contributing to the shuffled reconstruction, and calculated its proximity to the mean. We repeated this measure 1,000 times, and computed and empirical *p*-value as the proportion of true distances to the distribution mean that were smaller than the shuffled distances.

## 3. Results

### 3.1 Behavior

We order our findings by latency between training and test: the crab distribution on Day 8 with 0-day lag, the shell distribution on Day 8 with 7-day lag, the crab distribution on Day 21 with 13-day lag, and the shell distribution on Day 21 with 20-day lag.

At the shortest lag (0-day), null distributions revealed above chance proximal navigation to the mean for all ranges of rotation (360°: *p*_*rot*_=0.001; 180°: *p*_*rot*_=0.002; 90°: *p*_*rot*_=0.005). Navigation at the second shortest lag (7-day) was marginally more biased than a 360°rotated null (*p*_*rot*_=0.076), but not significantly different from the 180°rotated null (*p*_*rot*_=0.144) or 90°rotated null (*p*_*rot*_=0.301).

At the second-longest lag (13-day), test behavior again revealed navigation proximal to the mean as compared to the 360°(*p*_*rot*_=0.023) and 180°(*p*_*rot*_=0.044) null distributions, although only marginal in comparison to the 90°null (*p*_*rot*_=0.092). Proximity to the distribution mean at the longest lag (20-days) was significantly greater than the 360°null distribution (*p*_*rot*_=0.029), marginally greater than the 180°null (*p*_*rot*_=0.074), and not significantly different from the 90°null (*p*_*rot*_=0.138).

### 3.2 Reconstructed Location Representations

For the shell distribution, 46 information channels (out of 100) showed significant hippocampal place-coding activity. For the crab distribution, 17 information channels showed significant place-coding activity. We attribute the greater number of validated shell vs. crab channels to the patient exploring more of the arena at the beginning of the study, as she was gaining familiarity with the task and learning the first distribution. Critically, the number of validated information channels per distribution from training was held constant across test lags.

We investigated representational change from the shortest to the longest latency between training and test. Hippocampal activity patterns allowed us to reconstruct the distribution mean at the shortest lag (0-day: *p*_*rot*_=0.049) and second shortest lag (7-day: *p*_*rot*_ = 0.015). However, mPFC activity patterns did not contain information about the distribution mean at either of these lags (0-day: *p*_*rot*_=1; 7-day, *p*_*rot*_ = 0.333).

As hypothesized, we found the inverse at a longer latency. mPFC activity patterns allowed us to reconstruct the distribution mean at the second longest lag (13-day: *p*_*rot*_ = 0.009) and longest lag (20-day: *p*_*rot*_=0.048). However, hippocampal activity patterns did not contain information about the distribution mean at either of these lags (13-day: *p*_*rot*_=0.421; 20-day: *p*_*rot*_ = 0.539). These results indicate a double dissociation between the hippocampus and mPFC in the representation of spatial regularities as a function of time.

## 4. Discussion

We found evidence of statistical learning on two timescales by reconstructing place information from neural activity in the human brain. The learned spatial distributions switched from being represented only in the hippocampus at shorter latencies to being represented only in the mPFC at longer latencies.

Our findings are partially consistent with a previous study on the role of consolidation in accentuating stimulus overlap, which found that abstracted representations come online in mPFC only following a period of consolidation (Tompary & Davachi, 2017). However, that study additionally found parallel abstracted representations in the hippocampus after consolidation. The difference in length of consolidation may partly explain this discrepancy, with their study testing after one week of consolidation compared to our two and three week intervals. This additional time may have allowed for further transformation and weakening of the hippocampal representation. Given that the shell distribution was still represented in the hippocampus (but not mPFC) after one week, it may be that these two studies, with different tasks and stimuli, index different points along a trajectory of memory transformation. The endpoint of this trajectory in our study aligns well with previous rodent work showing cortically dependent spatial pattern representations emerging after 30 days of consolidation (Richards et al., 2014).

Our findings also serve to extend previous discoveries of ensemble-level spatial coding in the human brain. Single-unit recordings have identified sparse distributions of place cells (Ekstrom et al., 2003), and at the level of regional activity, multiple fMRI studies have revealed increasing hippocampal pattern similarity as a function of spatial distance between locations (Deuker, Bellmund, Schröder, & Doeller, 2016; Morgan, MacEvoy, Aguirre, & Epstein, 2011). Here, using inverted encoding models, we demonstrate more explicitly, and from fewer recordings, evidence of hippocampal place coding at the level of ensemble LFP. This result illustrates the high degree of information that can be derived from neural oscillations in a handful of macro electrodes, and provides a promising avenue for future navigation studies with iEEG.

This unique patient allowed us to track acquisition of spatial patterns across multiple timescales. An obvious limitation of the current study is the sample size of one. As a case study, our results should be interpreted with caution for their generalizability. However, this study provides initial support for the notion that consolidation operates over not only episodic memories, but extracted navigational patterns acquired online during learning. We additionally extend previous findings of the link between statistical learning and pattern similarity(Schapiro, Kustner, & Turk-Browne, 2012), arguing for an analogous process during human virtual navigation. Although it is unlikely that such a rare data collection opportunity will arise in a larger cohort of epilepsy patients, future noninvasive neuroimaging work could seek to verify the timeline discovered here. Our investigation provides a promising step in confirming the relationship between the hippocampus and mPFC in representing underlying structure in the world.

## 5. Funding

This work was supported by a National Institutes of Health (NIH) Grant R01MH069456 (to N. B. Turk-Browne), and by a CTSA Grant Number TL1 TR001864 from the National Center for Advancing Translational Science (NCATS), components of the NIH, and NIH roadmap for Medical Research (to K. N. Graves). Its contents are solely the responsibility of the authors and do not necessarily represent the official view of NIH.

---

**Fig. S 1.**
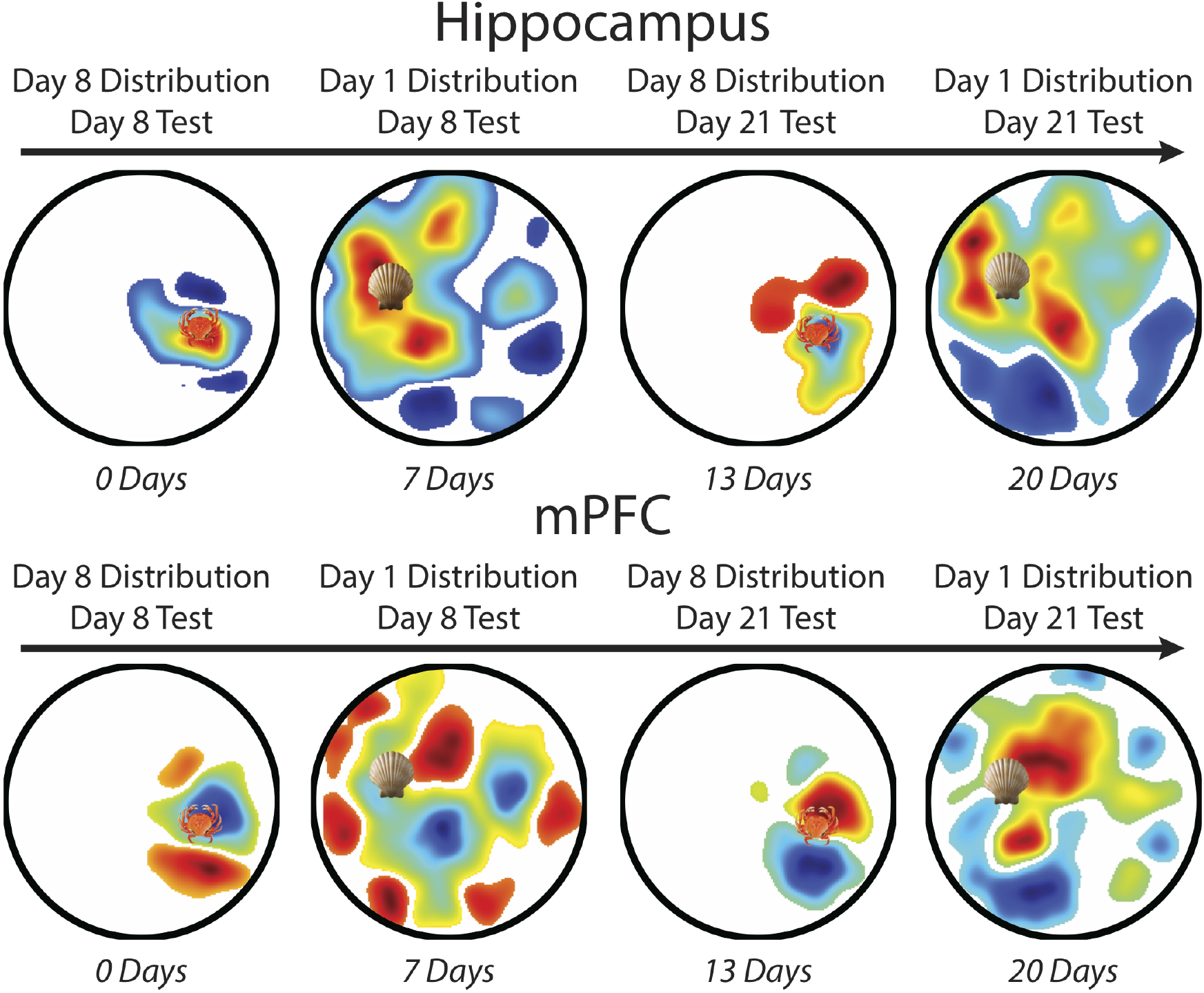
IEM reconstructions using a data threshold-based channel selection method, where channels were included in the model if the patient spent a cumulative minimum of 2500ms within that channel’s radius during search. Here, the Guassian shape parameters were manually set at 1 1/3 AU stimulus size and 1.2 AU receptive field size in order to accomplish a smooth reconstruction. The reconstruction from hippocampal data was significantly mean-proximal at both shorter latencies (0-days: *p*_*rot*_=0.009; 7-days: *p*_*rot*_=0.007) but not at the longer latencies (13-days: *p*_*rot*_=0.082; 20-days: *p*_*rot*_=0.546). The inverse was true for reconstructions from mPFC data, with significant mean-proximal reconstructions at the longer latencies (13-days: *p*_*rot*_=0.049; 20-days: *p*_*rot*_=0.002) but not the shorter latencies (0-days: *p*_*rot*_=0.613; 7-days: *p*_*rot*_=0.1).

## Notes

### Competing Interest Statement

The authors have declared no competing interest.

